# The humoral and cellular immune response of a third dose adenoviral-based vectored vaccine after a single or 2 shots of inactivated COVID-19 vaccine in healthy adults

**DOI:** 10.1101/2022.10.18.512323

**Authors:** Supaporn Suparak, Kobkaew Bumroongthai, Danai Jantapalaboon, Phanapat Pairoung, Sudarat Wongkidakarn, Chonlada Yodtub, Wiroj Puangtubtim, Warangluk Pimpapai, Sakulrat Soonthorncharttrawat, Panadda Dhepaksorn, Surakameth Mahasirimongkol

## Abstract

The coronavirus pandemic is a severe infectious respiratory disease which caused massive loss worldwide. Thailand was affected by the wild-type, and the Delta variant started in mid-2021. Due to the shortage of effective vaccines, the ministry of public health of Thailand suggested the heterologous vaccine scheme, which comprises inactivated COVID-19 vaccine (CoronaVac) and an adenoviral-based vectored vaccine (ChAdOx1). However, data on the humoral and cellular immune responses of the single or 2 shots of inactivated vaccine followed by the adenoviral-based vaccine are very limited. In this current study, the sera from participants who received either single or 2 shots of CoronaVac followed by the ChAdOx1 vaccine were evaluated for SARS-CoV-2 spike receptor-binding-domain (RBD) IgG. The cytokine level was also assessed using Luminex immunoassay. The PBMC were collected to evaluate spike-specific T-cell and B-cell responses. Participants who received 2 shots of CoronaVac followed by ChAdOx1 possessed significantly (*P*<0.0001) higher levels of spike RBD-specific IgG. They also exhibited a higher level of CD4+T-cell and IFN-gamma than those who received only 1 shot of CoronaVac followed by the ChAdOx1 vaccine. The volunteers who received two shots of CoronaVac followed by ChAdOx1 had a significantly (*p*<0.01) higher marginal B-cell response against wild-type SARS-CoV-2 S peptides than those who received only one shot of CoronaVac followed by ChAdOx1. Surprisingly, the class switch B-cell response to Delta variant SARS-CoV-2 S peptides of the volunteers who received 1 shot of CoronaVac followed by ChAdOx1 was significantly (*p*<0.01) higher than those who received 2 shots of CoronaVac. However, participants who received only a single shot of inactivated vaccine followed by an adenoviral-based vectored vaccine possessed a higher level of TNF-alpha and IL-6. This study indicated that boosting the ChAdOx1 as a third dose after completing 2 shots of CoronaVac induced strong humoral and cellular immune responses.

## Introduction

The coronavirus disease 2019 (COVID-19) pandemic, caused by the severe acute respiratory syndrome coronavirus 2 virus (SARS-CoV-2), first emerged in December 2019 [1]. Over 300 million confirmed cases of COVID-19 worldwide and over 5.5 million deaths in January 2022 [2]. Vaccination is one of the most effective strategies to reduce illness and mortality rates from the Covid-19 virus [3]. Various COVID-19 vaccines have been developed and used to fight against the pandemic [4]. However, as a developing country, Thailand faces significant challenges in acquiring and distributing adequate vaccine supplies [3]. The majority of Thai people could only access the first Covid-19 vaccine, which was inactivated COVID-19 from China, called CoronaVac^®^ (Sinovac), in mid-2021 [5]. The adenoviral-based vectored vaccine (ChAdOx1 AZD1222) was available in Thailand a few months after the first arrival of CoronaVac [6]. Due to the limited vaccine supplies, The Ministry of Public health of Thailand implemented a heterologous prime-boost vaccination strategy to create herd immunity to combat the COVID-19 pandemic [7].

Humoral and cell-mediated immune responses play essential parts in developing immunity after immunization with the vaccine against Covid-19 [8]. Most COVID-19 vaccines require at least 2 shots to evoke the humoral immune response, protecting and reducing the clinical symptom of COVID-19 [9, 10]. Two shots of inactivated vaccine elicit a higher CD4+ T-cell response to SAR-CoV-2 spike protein than only 1 dose of inactivated vaccine [11]. The previous report showed that participants who obtained 2 shots of CoronaVac followed by ChAdOx1 possessed a higher level of spike RBD-specific IgG and total immunoglobulins than those who received two-dose of CoronaVac or ChAdOx1 [5]. However, the data for SAR-CoV-2-specific cell-mediated immune response following vaccination with a heterologous prime-boost strategy are still scarce since most studies only focus on the humoral immune response. To fill the knowledge gap, we characterized various parameters of SARS-CoV-2-specific cellular and humoral immune responses induced by once or twice CoronaVac followed by ChAdOx1 in healthy Thai individuals.

## Materials and Methods

### 1. Study design and participants

All experiments with humans were conducted in accordance with the ethical approval by the ethical committee of the Department of Medical Sciences on 23^rd^ July 2021 with approval number; MOPH 0625/EC095 and study code; 15/2564. All participants agreed and signed the consent forms before blood donation. Three millilitres of blood samples were taken from donors by venipuncture into the K2EDTA vacutainer tube (Becton, Dickinson Limited (BD). Blood samples were collected at baseline before vaccination (pre-booster) and 2 weeks and 4 weeks intervals after vaccination the third dose.

### 2. PBMC isolation

Peripheral blood mononuclear cells (PBMCs) were isolated within 4-6 hours after blood collection by density gradient centrifugation. First, blood samples were diluted with an equal amount of phosphate buffer saline solution (PBS) and slowly layered on top of the same volume of Lymphoprep density gradient medium (Stemcell Technologies Inc.). After that, the solutions were centrifuged at 800 g for 10 minutes at 18°C. PBMCs were transferred into 15 mL conical centrifugation tubes and washed twice with 2% albumin in PBS. The PBMCs were resuspended in freezing solution (10%DMSO in FBS) and kept at −80°C until use.

### 3. Quantification of the lymphocyte population

Lymphocyte subsets were analyzed using cyto-Stat tetra CHROME (CD3/ CD4/CD8/CD45) reagent (Beckman Coulter, Switzerland). The analysis was performed according to the product manual. Fifty microliters of EDTA anticoagulated blood were mixed with 10 mL of antibody solution and incubated for 15 minutes in the dark at 4°C.

After that, the cell suspensions were washed and lysed with 450 microlitres of BD FACS lysing solution (BD Biosciences, USA) and incubated for another 15 minutes at room temperature. The samples were analyzed using BD FACs Canto II flow cytometer (BD Biosciences, San Jose, USA).

### 4. The measurement of anti-SAR-CoV-2 RBD IgG level

The level of anti-RBD IgG antibodies was measured in human plasma by chemiluminescent microparticle assay (CMIA) using the SARS-CoV-2 IgG II Quant (Abbott, USA) on the ARCHITECT I System (Abbott, Abbott Park, USA). The plasma samples were collected and stored at −80°C until used. The antibody concentration ranged from 6.8 Abbott Arbitrary Units per milliliter (AU/mL) to 40,000 AU/mL. Antibody levels greater than 50 AU/mL were deemed as seropositive.

### 5. Peptides for the T-cell stimulation assay

The spike – S1 peptide pools of Wuhan wild-type SAR-CoV2 were purchased from Genscript (cat # RP30027). The Peptivator peptide pools for the variant of concerns and B. 1.617 (delta), were purchased from Miltenyi (USA). The Wuhan wild-type S peptides contain 158 and 157 peptides. The delta strain peptide pools contain 157 & 158 peptides, 15mers with 11 amino acids overlapping the full-length of spike glycoprotein of SAR-CoV2 B.1.617. All lyophilized peptides were generated at greater than 95% purity. The wild-type strain was reconstituted with DMSO and diluted with PBS to get the less than 1% of DMSO in the ready-to-use stock solution. The variant strains were reconstituted with PBS. All of the reconstituted peptides were kept at −20°C until used.

### 6. T-cell restimulation with SAR-CoV2 peptide

Cryotubes containing frozen PBMCs were placed at 37°C in a bath until thawed cells. Cells were pipetted into 15 mL tubes and added 3 ml of warm RPMI-1640 media. Then, cells were washed 2 times by centrifugation at 350×g for 5 min. After that, cells were resuspended in complete RPMI-1640 supplemented with 5% human platelet lysate and 1% streptomycin-penicillin. Cell viability was checked with Trypan Blue. For stimulation, 2.5×10^5^ cells were diluted with an equal volume of each of the diluted spike protein peptide pools. The wild-type peptide pools were prepared in the complete RPMI medium at the final concentration of 2 μg/mL. The variant strains were prepared in sterile PBS at the final concentration of 4 UI. The PBMCs were incubated with Dyna bead (Thermofisher, USA) as positive control and a complete RPMI medium as the negative control. Then, all the PBMCs were incubated for 16-18 hours at 37°C with 5% CO_2_. After incubation, cells were centrifuged at 350g for 5 minutes and collected for the intracellular cytokine staining assay. The supernatants were transferred into the Eppendorf and stored at −80°C for cytokine quantification by Multiplex Immunoassay.

### 7. Intracellular cytokine staining

The stimulated PBMCs were washed twice with PBS pH7.2 buffer and then permeabilized using the PerFix-nc^®^ reagent kit (Beckman Coulter, USA). Briefly, 25 μL of R1 buffer was added to 50 μl of washed cell and incubated for 15 minutes at room temperature. Then, the cell suspension was washed once with PBS pH7.2. After that, 25 μL of fetal bovine serum and 300 μl R2 Buffer were added to the cell suspension and transferred to DURAClone IF T Activation Tube (Beckman Coulter, USA), incubated in the dark for 45 minutes. Cells were washed and resuspension with 500 μL of diluted R3 solution. All data were acquired using the CytoExpert Software (Beckman Coulter, USA) (version 2.0).

### 8. IFN-gamma and IL-2 ELISpot assay

The ELISpot assay was performed using PVDF membrane 96-well microplates (R&D Systems) following the manufacturer’s guidelines. The isolated PBMCs were thawed and rested in RPMI media for 1 hour. After that, 200 μl of 2 x 10^6^ PBMC/mL cell suspension (2.5 x 10^5^ PBMC per well) were stimulated for 16-18 hours with either RPMI media (negative control), Dyna bead (positive control), each SAR-CoV-2 peptides mixes (2 μg/mL of either wild-type or Delta variant SAR CoV-2 S peptides. Each sample was performed in duplicate and then incubated at 37°C with 5% CO_2_. Cell viability of more than 80% was used in this assay.

The number of SARS-CoV-2 specific IFN-gamma and IL-2-secreted T-cells per 2.5 x 10^5^ PBMCs were determined using *ImmunoSpot S6 Ultimate* Analyzer (CTL, Cleveland, USA) and presented as spot forming unit per 2.5 hundred thousand PBMCs (SFU/2.5×10^5^).

### 9. Cytokine quantification by Multiplex Immunoassay

Cytokine production was determined by Luminex technology. The supernatants obtained from the T-cell stimulation assay were collected and stored at −80°C until analysis. Luminex 200 was used to analyze the results according to the manufacturer’s instructions. A Th1/Th2 cytokine panel (ProcartaPlex™Panel, Thermo Fisher Scientific, Catalog number: EPX060-10009-901) was used to detect cytokines, including IFN-gamma, IL-12p70, IL-4, IL-6, and TNF-alpha. Briefly, 50 μL of undiluted supernatant was added to the antibody magnetic beads in each of the 96-well plate wells for 60-120 minutes on a plate shaker at room temperature. Next, detection antibodies were added to each well and incubated for 30 minutes on a plate shaker, followed by streptavidin phycoerythrin incubation (50 μL per well). After 30 min of incubation, the beads-sample mixtures were resuspended in 25 μL of reading Buffer. The cytokine expression profile was characterized with Luminex 200™ instrument (Luminex Corporation, Austin, TX, USA) following the manufacturer’s instructions. The concentrations of cytokines were calculated by using xPONENT^®^ version 3.1 software.

### 10. Statistical analysis

Statistical analysis was calculated using GraphPad Prism 8 software (GraphPad Software Inc., San Diego, CA, USA). The results from the 2 groups were determined using either student *t*-test or the Mann-Whitney test. The multiple groups were tested with ANOVA multiple comparison test. A value of *P* ≤ 0.05 was considered statistically significant.

## Results

### 1. The anti-RBD-specific IgG antibody response

To evaluate the circulating IgG-specific RBD of the SARS-CoV2 antibody, the serums from volunteers were collected 4 weeks after immunization with ChAdOx1 and measured the antibody level using CMIA. The average IgG antibody of the participants who received either 1 or 2 shots of CoronaVac followed by the ChAdOx1 vaccine was 5,077 AU/mL and 14,008 AU/mL, respectively. Interestingly, the results showed that the level of IgG-specific RBD of SARS-CoV2 antibody of participants who received 2 shots of CoronaVac followed by ChAdOx1 vaccine was significant (*P*<0.0001) higher than participants who received only 1 dose of CoronaVac before ChAdOx1 vaccination as seen in **Fig 1.**

**Fig 1.**
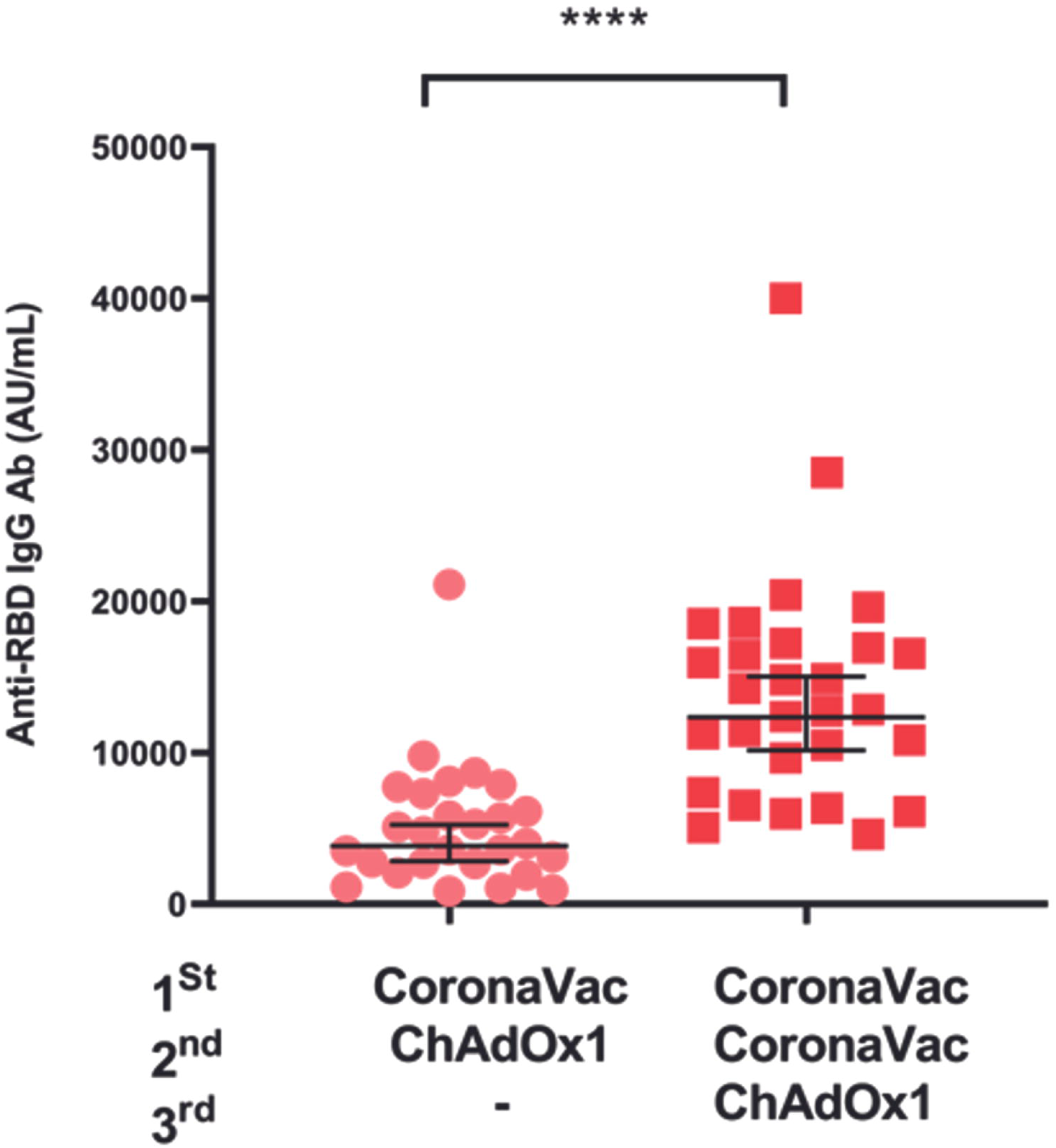
Antibody responses against SAR-CoV2-RBD following ChAdOx1 Vaccination after 1 or 2 CoronaVac shots. The anti-SAR-CoV2 RBD IgG antibody level was compared between volunteers who received either 1 dose (n = 27) or 2 doses (n = 29) of CoronaVac followed by ChAdOx1 at 4 weeks after immunization. Statistical analysis was performed using the student *t*-test. \*\*\*\**P*<0.0001.

### 2. T-cell response to spike protein-peptide pools in wild-type strains and variants of concern

The human IFN-gamma/IL-2 Dual-Color ELISpot Kits evaluated the spike-specific T-cell response. The responses of T-cells from volunteers who received 1 CoronaVac followed by ChAdOx1 Vaccine (n=12) were compared to the volunteers who received 2 shots of CoronaVac followed by ChAdOx1 Vaccine (n=12). There was no significant difference in IFN-gamma and IL-2 levels against wild-type and VOC following the immunization with the 2nd or third dose with ChAdOx1, as shown in **Fig 2**.

**Fig 2.**
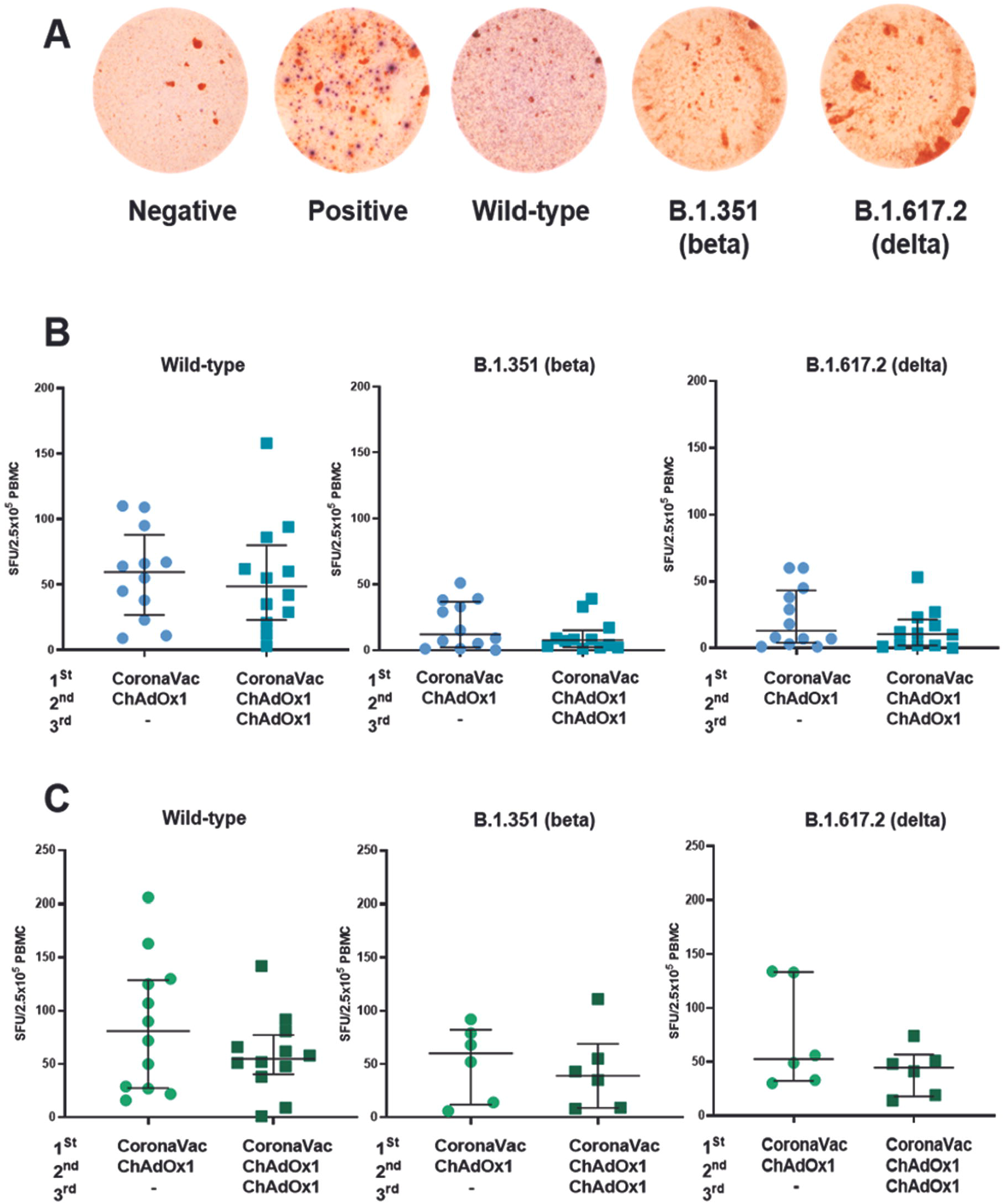
Spike-specific T-cell response following ChAdOx1 Vaccine after 1 or 2 CoronaVac shots evaluated by T-cell ELISpot. (A) Representative images of the ELISpot wells against wild-type and variant peptides. IFN-gamma (blue spots) and IL-2 (red spots) were secreted from T-cell stimulated with pools of Spike peptides derived from SARS-CoV-2 WT and variants of concern. (B) INF-gamma ELISpot and (C) IL-2 results assess following the second or third vaccination with the ChadOX1 vaccine. Cryopreserved PBMCs were stimulated with S1 spike peptides pools of SAR-CoV-2 Wild type and variants of concern to determine the specific responses of T cells. The results were expressed as spot forming units per 2.5 hundred thousand PBMCs (SFU/2.5×10^5^). The responses of T-cells from volunteers who received 1 CoronaVac followed by ChAdOx1 Vaccine (n=12) were compared to the volunteers who received 2 doses of CoronaVac followed by ChAdOx1 Vaccine (n=12). Statistical analysis was determined by Mann-Whitney test.

### 3. Immunophenotyping of TBNK subpopulations from vaccinated volunteer

The immunostaining analysis of CD4+T-cell, CD8+T-cell, B-cell, and NK cell from the whole blood samples were analyzed in volunteers who received 1 CoronaVac followed by ChAdOx1 (n = 30) with the volunteers who received 2 doses of CoronaVac followed by ChAdOx1 (n = 20). The amount of CD4+T-cell from volunteers who received 2 doses of CoronaVac followed by ChAdOx1 Vaccine was significantly (*p*<0.0001) higher than the volunteers who received only 1 shot of CoronaVac followed by ChAdOx1 Vaccine. On the other hand, the level of CD4+T-cell from volunteers who received 2 doses of CoronaVac followed by ChAdOx1 Vaccine was significantly (*p*<0.0001) lower than the volunteers who received only 1 dose of CoronaVac followed by ChAdOx1 Vaccine. There was no significant difference in the amount of B-cell and NK cells between both groups, as shown in **Fig 3.**

**Fig 3.**
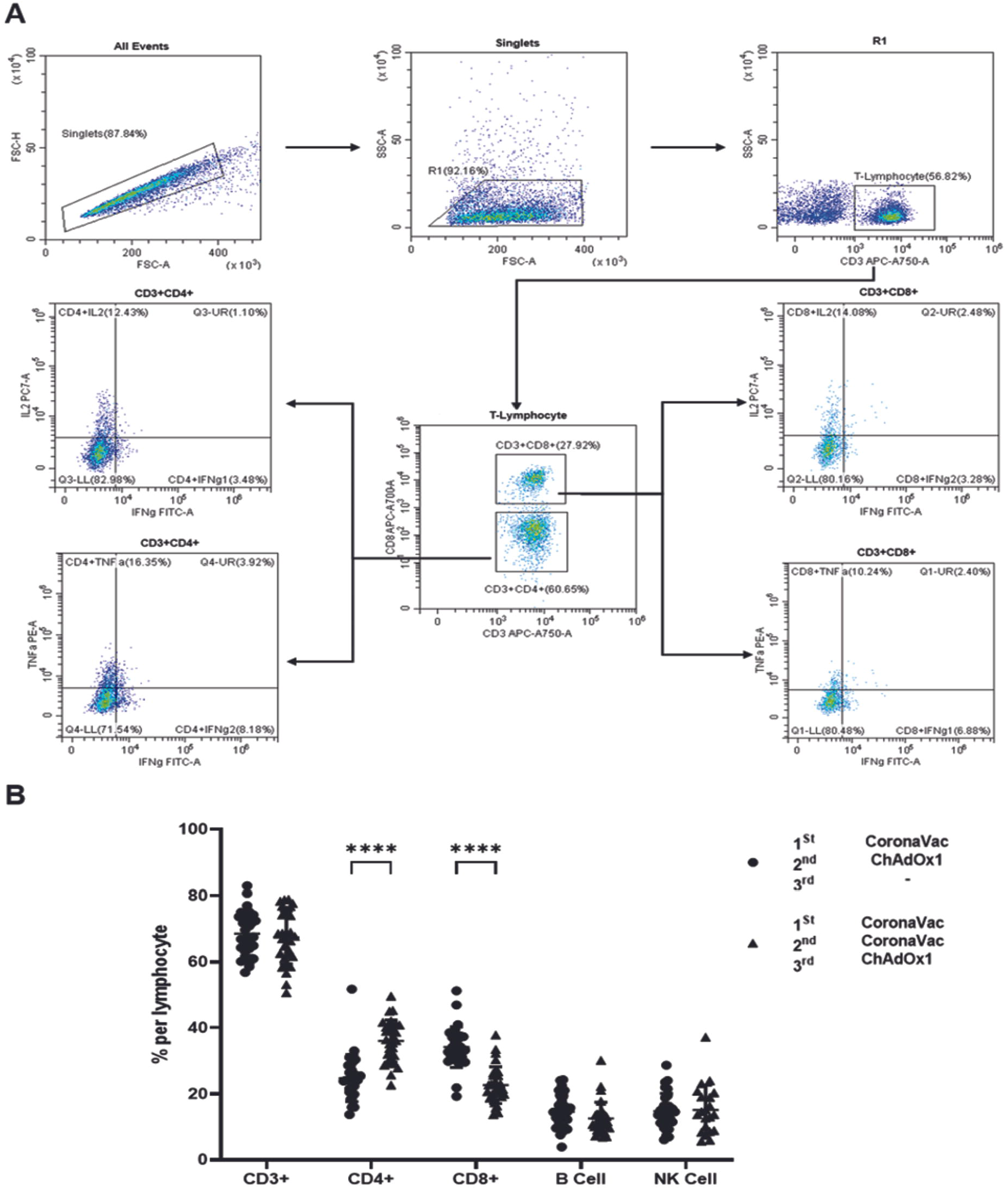
Immunophenotyping of T-cells, B-cells, and NK cells subpopulations from whole blood of volunteer vaccinated with either 1 shot or 2 shots of CoronaVac followed by ChAdOx1 vaccine. (A) The gating strategy for CD3+T-cells detection using multiparameter flow cytometry. (B) Flow cytometric analysis showed CD4+T-cells, CD8+T-cells, B-cells, and NK-cells characterization compared between 1 dose (n = 30) or 2 doses (n = 20) of CoronaVac followed by ChAdOx1. Statistical analysis was performed using ANOVA multiple comparison test. \*\*\*\**P*<0.0001

### 4. Cytokine quantification assays

This experiment tested cytokine quantification of Th1/Th2 using 6 cytokine types, including IFN-gamma, TNF-alpha, IL-4, IL-6, and IL-12p70, in response to T-cell stimulated with pools of Spike peptides derived from SARS-CoV-2 wild type and Delta variants of concern. The cytokine quantification was compared in volunteers who received only 1 shot of CoronaVac vaccine followed by ChAdOx1 Vaccine (n=11) with the volunteers who received 2 doses of CoronaVac vaccine followed by ChAdOx1 vaccine (n=11). The result was statistically significant in IL-6 and IL-12p70 (**Fig 4**).

**Fig 4.**
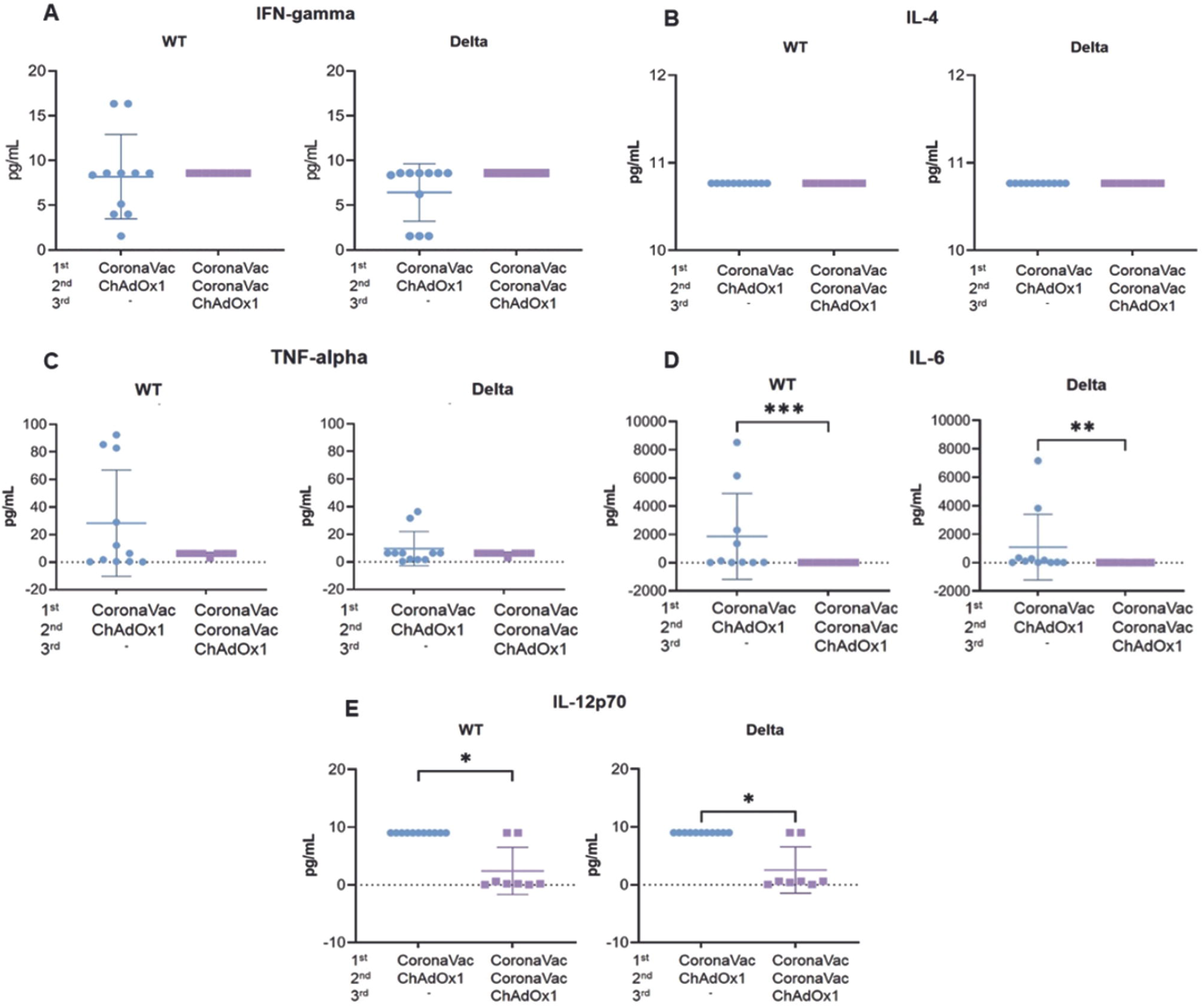
Cytokine quantification assays. Comparison of (A) IFN-gamma, (B) IL-4, (C) TNF-alpha, (D) IL-6, and (E) IL-12p70 cytokine responses to wild-type (n =11) and Delta variant (n =11) SARS-CoV-2 S peptides of volunteers vaccinated with either 1 or 2 CoronaVac shots followed by ChAdOx1 vaccine. Each dot represents of cytokine concentration individual sample. The statistical analysis was performed using the Mann-Whitney test. \**p* < 0.05, \*\**p* < 0.001, and \*\*\**p* <0.0001.

### 5. Cellular immune response against wild-type and Delta (B.1.617.2) variant SARS-CoV-2 S Peptides

Isolated PBMC from healthcare volunteers immunized with different vaccines were stimulated with 5 μM of wild-type and Delta variant SARS-CoV-2 S peptides after 4 weeks of vaccination. Intracellular TNF-alpha response to wild-type SARS-CoV-2 S peptides of the volunteers who received only 1 shot of CoronaVac followed by ChAdOx1 was significantly (*p*<0.01) higher than in the group that received 2 shots of CoronaVac followed by ChAdOx1. When stimulated with Delta variant SARS-CoV-2 S peptides, volunteers who received 1 dose of CoronaVac had significantly (*p*<0.01) higher TNF-alpha expression levels than those who received only 2 shots of CoronaVac followed by ChAdOx1. Surprisingly, the expression of INF-gamma in volunteers who received two doses of CoronaVac followed by ChAdOx1 was significantly (*p*<0.001) increased in response to wild-type SARS-CoV-2 S peptides. The results were similar to those stimulated with Delta SARS-CoV-2 S peptides. However, there was no significant difference in intracellular IL-2 expression between 2 groups of people who received different types of vaccines stimulated with wild-type and Delta SARS-CoV-2 S peptides, as shown in **Fig 5**.

**Fig 5.**
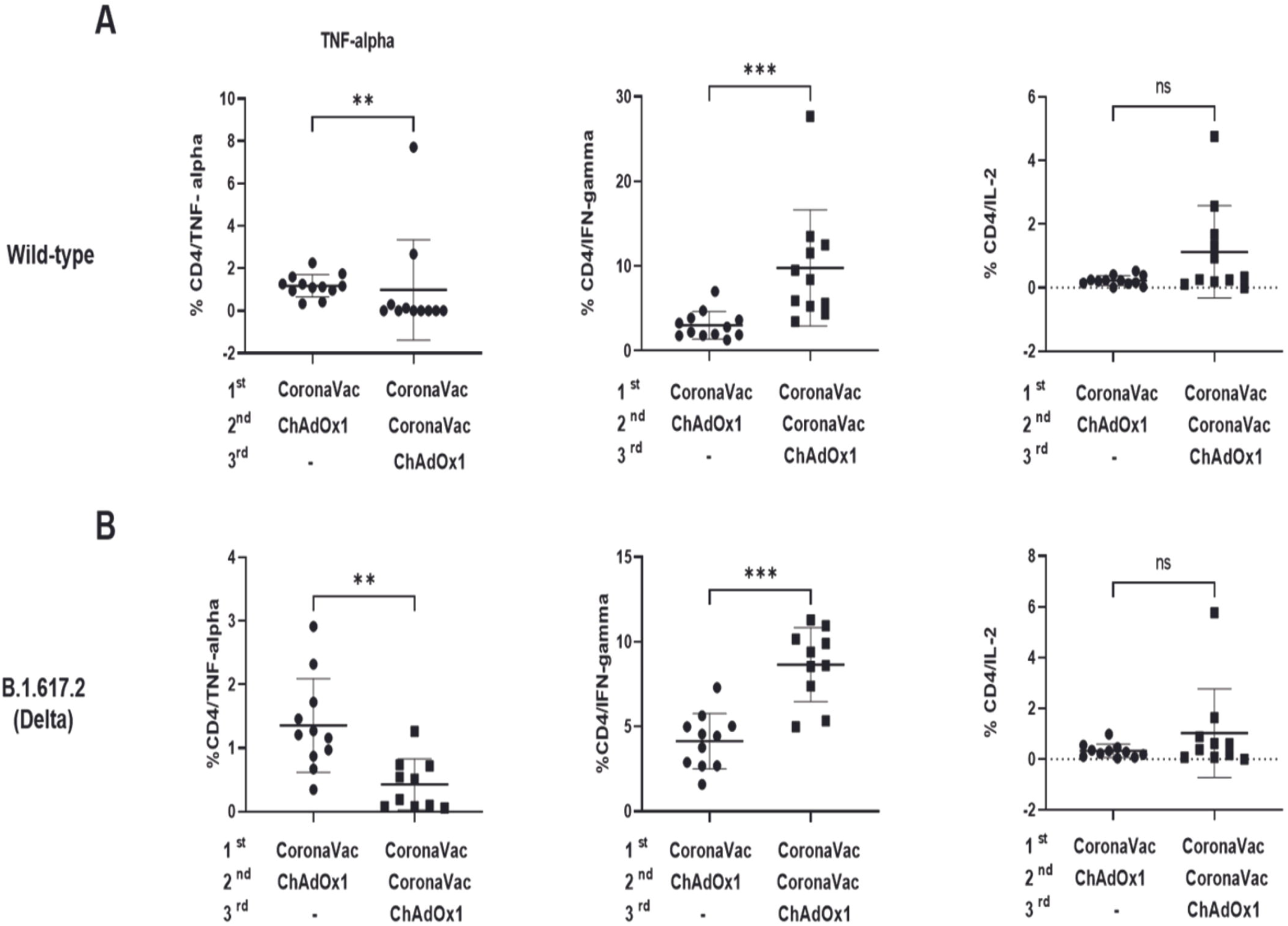
Cellular immune responses against wild-type and Delta (B.1.617.2) variant SARS-CoV-2 S peptides following ChAdOx1 vaccination after 1 or 2 CoronaVac shots. Intracellular immunostaining showed T-cell response through cytokine expression level after being stimulated with wild-type (A) and Delta variant SARS-CoV-2 S Peptides (B) compared between 1 dose (n = 12) or 2 doses (n = 10) of CoronaVac followed by ChAdOx1 at 4 weeks after immunization. Statistical analysis was performed using the Mann-Whitney test. \*\**P*<0.01, \*\*\**P*<0.001.

B-Cells responses were observed using stimulated PBMC with peptide variants as described previously. As shown in **Fig 6**, the marginal B-cell response to wild-type SARS-CoV-2 S peptides of the volunteers who received 2 shots of CoronaVac followed by ChAdOx1 was significantly (*p*<0.01) higher than those who received only 1 shot of CoronaVac followed by ChAdOx1. The responses of transitional B-cell and class switch B-cell against wild-type SARS-CoV-2 S peptides were higher in volunteers who received only one shot of CoronaVac followed by ChAdOx1, but this was not statistically significant. Surprisingly, when stimulated with Delta variant SARS-CoV-2 S peptides, the class switch B-cell of the volunteers who received 2 shots of CoronaVac followed by ChAdOx1 was significantly (*p*<0.01) lower than those who received only 1 shot of CoronaVac followed by ChAdOx1. **Fig 7** shows no significant difference in Class-unswitched memory B-cells (CD19+/IgM+/IgD+/CD38-/CD27+) responses to wild-type and Delta SARS-CoV-2 S peptides between two groups of people who received different types of vaccines.

**Fig 6.**
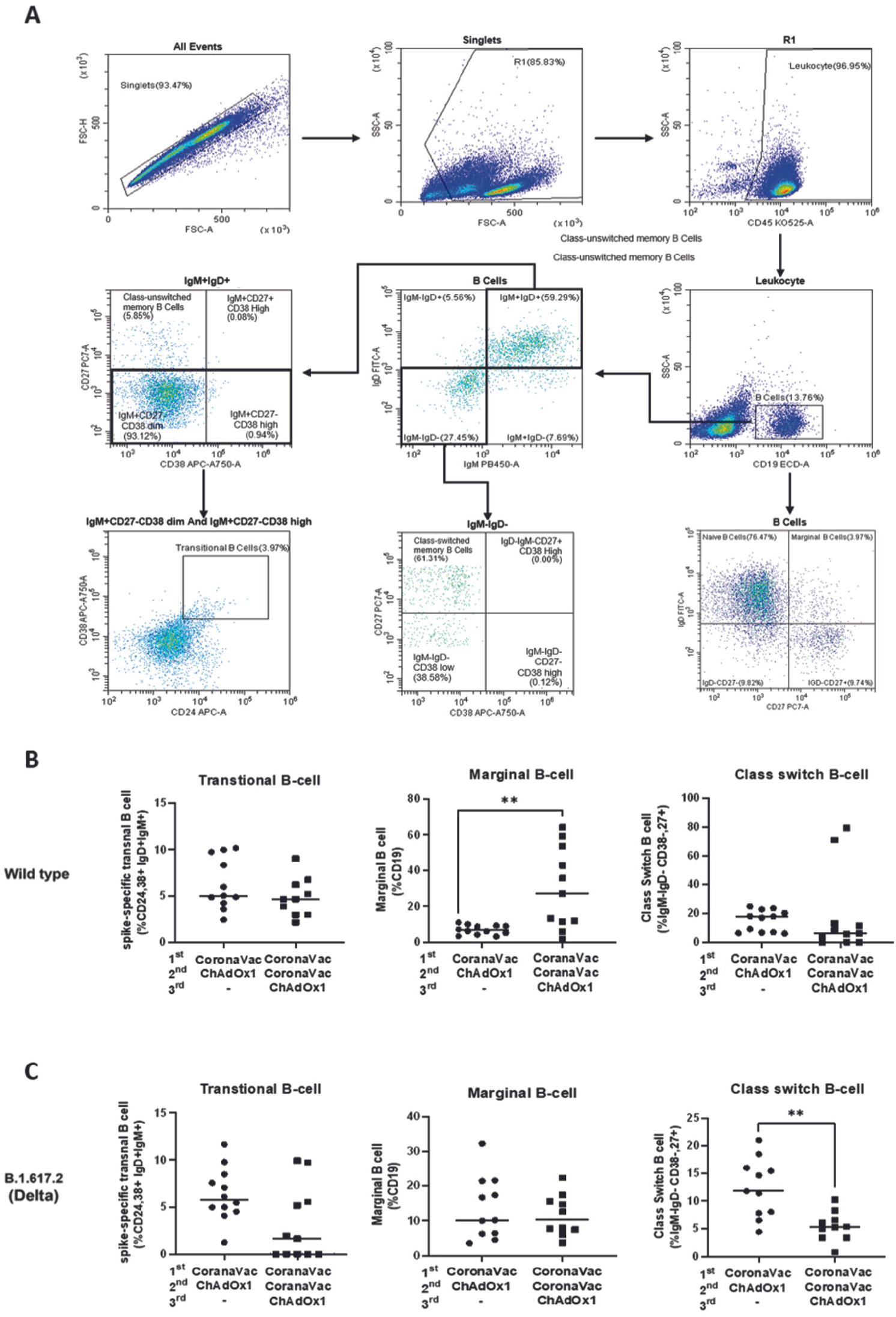
The flow cytometric analysis for B-cell responses against the wild-type and Delta variant SARS-CoV-2 S peptides. (A) The gating strategy for spike-specific B-cells detection using multiparameter flow cytometry. (B) The specific B-cell responses to wild-type SARS-CoV-2 S peptides of volunteers immunized with either 1 shot (n=11) or 2 shots (n=10) of CoronaVac followed by the ChAdOx1 vaccine. (C) The specific B-cell responses against Delta variant SARS-CoV-2 S peptides of volunteers vaccinated with either 1 shot (n=11) or 2 shots (n=10) of CoronaVac followed by ChAdOx1 vaccine. The statistical analysis was performed using the Mann-Whitney test. \*\**P*<0.01.

**Fig 7.**
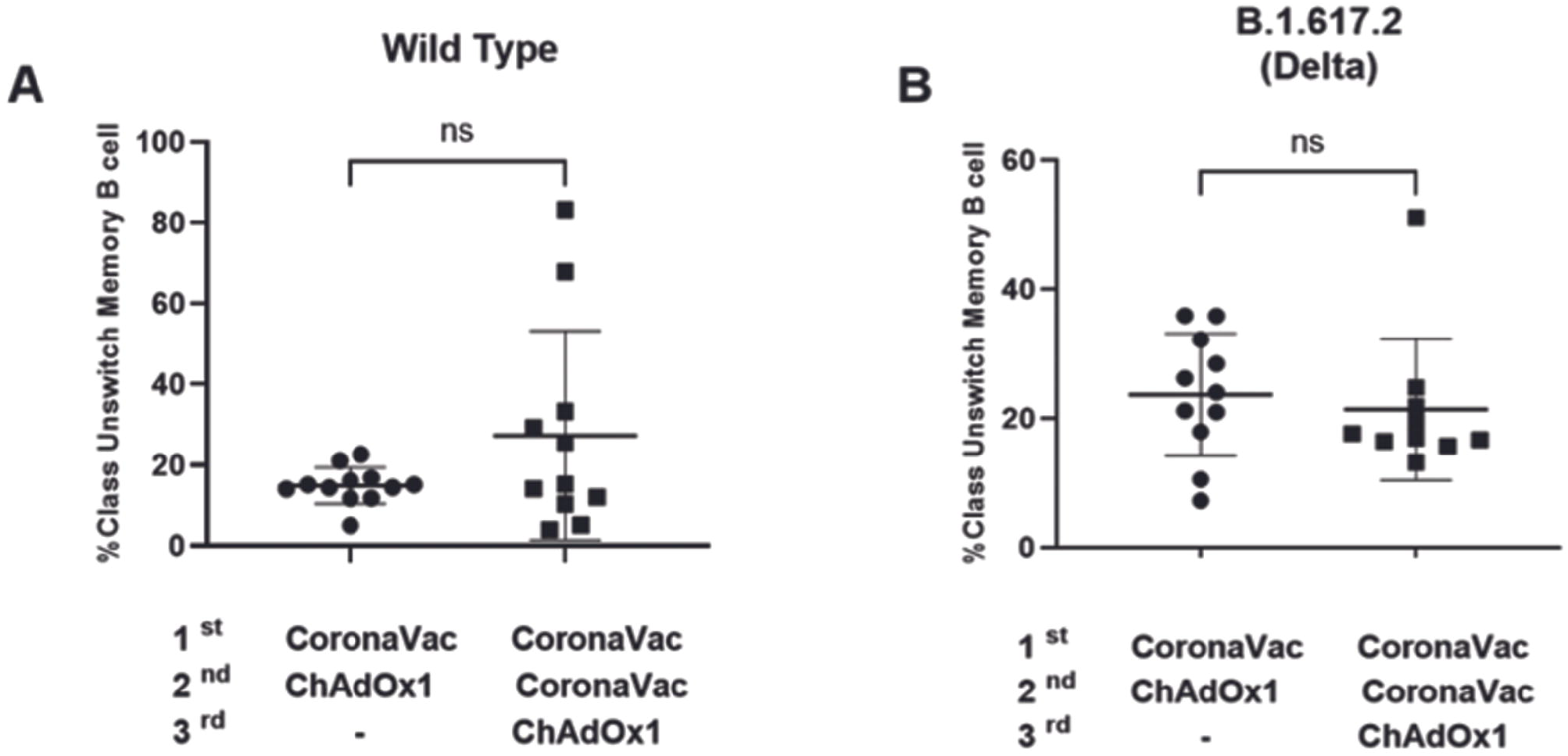
B-cell responses to wild-type and Delta variant SARS-CoV-2 S peptides of volunteers vaccinated with either 1 or 2 CoronaVac shots followed by ChAdOx1 vaccine. Stimulated PBMC Immunostaining analysis showed B-cell response through Class unswitched memory T-cell (CD19+/IgM+/IgD+/CD38-/CD27+) characterization after stimulated with wild-type (A) and Delta variant (B) SARS-CoV-2 S peptides compared between 1 dose (n = 12) or 2 shots (n = 11) of CoronaVac followed by ChAdOx1 at 4 weeks after immunization. Statistical analysis was performed using the Mann-Whitney test. ns, non-significant.

## Discussion

The SAR-CoV-2 pandemic crisis is still ongoing in several parts of the world. The mass vaccine is one of the best strategies to encounter this problem to create herd immunity against SAR-CoV-2 infection [12]. The heterologous vaccination is currently considered in developing countries due to the shortage of vaccines. In this study, we investigated the IgG antibody response against receptor-binding domain (RBD) of S1 spike protein of SARS-CoV-2 and cellular immune response following the 1 or 2 doses of inactivated SAR-CoV-2 vaccine CoronaVac followed by the adenoviral-based vectored vaccine (ChAdOx1 AZD1222). We demonstrated the antibody level against the SAR-CoV-2 induced by either 1 or 2 shots of CoronaVac followed by ChAdOx1 at 4 weeks after immunization. The antibody response induced by 2 doses of CoronaVac followed by ChAdOx1 was greater up to almost 3-fold compared to only 1 dose of CoronaVac followed by ChAdOx1 vaccination. These data suggested that a third dose of the COVID-19 vaccine elicits greater immunogenicity than 2 doses. Our results were similar to previous research, which indicated that boosting ChAdOx1 after completing 2 shots of CoronaVac elicited a higher number of RBD-specific IgG than those without the boosting [5]. Several studies found that three doses of inactivated or mRNA vaccine resulted in a more significant immunogenic response than two doses [13, 14].

Cellular immune responses were evaluated in the participants who received a booster dose of ChAdOx1 following either 1 or 2 shots of CoronaVac 4 weeks after the third vaccination. We demonstrated that vaccinees who received 1 shot of ChAdOx1 after 2 shots of CoronaVac exhibited a higher level of non-specific CD4+ T-cell than those who received 1 shot of ChAdOx1 after only 1 dose of CoronaVac. Unlike the mRNA vaccine, the inactivated vaccine contains the entire viral antigen and the immunogenic N protein, not just the spike protein [15]. The whole viral antigen induces T-cell immunity, which may explain this study’s high CD4+ T-cell response. The intracellular cytokine immunostaining technique also measured the specific-cellular immune response for wild-type SAR CoV-2 and Delta variant peptide protein. We discovered that those who received three doses of heterologous vaccine produced more IFN-gamma against wild-type SAR CoV-2 and Delta variant than those who received two doses. Adenoviral vector vaccines have been demonstrated to induce the aggressive cellular immune response and drive Th1 expansion [16]. The response against the spike protein may be influenced by the strong Th1 reaction to the adenoviral vector in the live vaccine. However, participants who received a single dose of CoronaVac followed by ChAOx1 demonstrated higher T-cell intracellular TNF-alpha against wild-type SAR CoV-2 and Delta variant than those who received 2 doses of CoronaVac followed by ChAOx1. The previous study showed similar results: boosting with CoronaVac induced the activation of CD4+T-cell and secretion of IFN-gamma against wild-type SAR-CoV-2, Delta, and Omicron variants [17]. Goa and colleagues demonstrated that CD4+T-cell response against wild-type strain is cross-reactive against Omicron strain in participants who received BNT162b2 mRNA vaccine [18].

The cytokines secretion (IFN-gamma, TNF-alpha, IL-4, IL-6, and IL-12p70, respectively) in response to T-cell stimulated with pools of Spike peptides derived from SARS-CoV-2 wild type and Delta variant were also analyzed by Luminex immunoassay. Surprisingly, IL-6 and IL-12p70 were significantly higher in participants immunized by a single shot of CoronaVac followed by ChAdOx1 compared to those who were immunized with 2 doses of CoronaVac 4 weeks after vaccination. Many types of vaccines increase IL-6, such as influenza and the Bacillus Calmette-Guérin vaccine [19]. For the influenza vaccine, the natural killer cell produces a high level of interferon-gamma, which stimulates CD11b+ dendritic cells to release IL-6 [20]. A recent study has reported the response of NK-cell followed by ChAdOx1, which may explain and support the increase of IL-6 in this current study [21].

In humoral immune response to vaccination, B-cells are stimulated by antigens in vaccines and developed into either plasma cells or memory B cells [22]. Memory B-cells are essential in vaccine-induced immunity since they rapidly produce antibodies to respond and protect against pathogen invasion [23]. The class switch B-cell is responsible for switching IgM production to IgA or IgG production [24]. Surprisingly, after weeks of vaccination with 2 shots of CoronaVac followed by the ChAdOx1 vaccine, the percentage of class switch B-cells was significantly lower than after only 1 shot of CoronaVac followed by the ChAdOx1 vaccine. Germinal centers in secondary lymphoid organs serve as microenvironments for the affinity development of humoral immunity 2 weeks after vaccination [25]. After T-cell-dependent activation, B-cells migrate to germinal centers, specifically differentiated into high-affinity memory B-cells and plasma cells [25]. This theory is supported by our data demonstrating that the inactivated vaccination can promote humoral immunity, reflected in the marked decline in B-cell counts. The recent study also demonstrated a similar result in the substantially reduced humoral B-cell levels measured using the CYTOF method in humans after a month of immunizing with the inactivated vaccine [26]. The specific SAR-CoV 2 antibody level was maintained for at least 6 months after vaccination with the inactivated vaccine [27]. The high antibody level after the vaccination highlights the memory immune response and signifies the importance of booster immunization.

The current study has some limitations. We included a small number of participants who received either 1 or 2 doses of inactivated vaccine followed by 1 shot of adenoviral-based vaccine. Second, we did not measure the baseline IgG antibody response against the receptor-binding domain (RBD) of S1 spike protein SARS-CoV-2 before and after the first vaccination. In addition, the samples for T-cell and B-cell responses were low due to this study’s limited time and budget. Furthermore, we only included Wild type SAR CoV-2 and Delta variants because they were the dominant strains during the study. The investigation of vaccine protection against emerging variants should be considered.

In conclusion, boosting the ChAdOx1 as a third dose after completing 2 doses of CoronaVac induced strong both humoral and cellular immune responses.

## Ethics Statement

The ethical approval of this study was approved by the ethical committee of the Department of Medical Sciences with approval number; MOPH 0625/EC060, study code; 15/2564, and date of approval; 23^rd^ July 2021.

## Acknowledgement

The authors would like to thank the Clinical Research Centre (CRC) from the Department of Medical Sciences and all the participants for supporting this project.

## References

1. Zheng J. SARS-CoV-2: an Emerging Coronavirus that Causes a Global Threat. International journal of biological sciences. 2020;16(10):1678–85. PubMed PMID: 32226285. Pubmed Central PMCID: 7098030.

2. Leigh JP, Moss SJ, White TM, Picchio CA, Rabin KH, Ratzan SC, et al. Factors affecting COVID-19 vaccine hesitancy among healthcare providers in 23 countries. Vaccine. 2022 Jul 29;40(31):4081–9. PubMed PMID: 35654620. Pubmed Central PMCID: 9068669.

3. Li J, Song M, Guo D, Yi Y. Safety and Considerations of the COVID-19 Vaccine Massive Deployment. Virologica Sinica. 2021 Oct;36(5):1097–103. PubMed PMID: 34061319. Pubmed Central PMCID: 8167387.

4. Kudlay D, Svistunov A. COVID-19 Vaccines: An Overview of Different Platforms. Bioengineering. 2022 Feb 12;9(2). PubMed PMID: 35200425. Pubmed Central PMCID: 8869214.

5. Yorsaeng R, Suntronwong N, Phowatthanasathian H, Assawakosri S, Kanokudom S, Thongmee T, et al. Immunogenicity of a third dose viral-vectored COVID-19 vaccine after receiving two-dose inactivated vaccines in healthy adults. Vaccine. 2022 Jan 24;40(3):524–30. PubMed PMID: 34893344. Pubmed Central PMCID: 8639402.

6. Pinpathomrat N, Intapiboon P, Seepathomnarong P, Ongarj J, Sophonmanee R, Hengprakop J, et al. Immunogenicity and safety of an intradermal ChAdOx1 nCoV-19 boost in a healthy population. NPJ vaccines. 2022 May 13;7(1):52. PubMed PMID: 35562372. Pubmed Central PMCID: 9106663.

7. Watcharananan SA, Nadee C, Kongsuwattanaleart P, Sangthong N, Ngorsakun P, Vimonvattaravetee P, et al. Rates, types, and associated factors of acute adverse effects after the first dose of ChAdOx1 nCoV-19 vaccine administration in Thailand. IJID Regions. 2022 Mar;2:35–9. PubMed PMID: 35721432. Pubmed Central PMCID: 8626869.

8. Gadani SP, Reyes-Mantilla M, Jank L, Harris S, Douglas M, Smith MD, et al. Discordant humoral and T cell immune responses to SARS-CoV-2 vaccination in people with multiple sclerosis on anti-CD20 therapy. EBioMedicine. 2021 Nov;73:103636. PubMed PMID: 34666226. Pubmed Central PMCID: 8520057.

9. Al Kaabi N, Zhang Y, Xia S, Yang Y, Al Qahtani MM, Abdulrazzaq N, et al. Effect of 2 Inactivated SARS-CoV-2 Vaccines on Symptomatic COVID-19 Infection in Adults: A Randomized Clinical Trial. Jama. 2021 Jul 6;326(1):35–45. PubMed PMID: 34037666. Pubmed Central PMCID: 8156175.

10. Zhang R, Khong KW, Leung KY, Liu D, Fan Y, Lu L, et al. Antibody Response of BNT162b2 and CoronaVac Platforms in Recovered Individuals Previously Infected by COVID-19 against SARS-CoV-2 Wild Type and Delta Variant. Vaccines. 2021 Dec 7;9(12). PubMed PMID: 34960189. Pubmed Central PMCID: 8705363.

11. Li Z, Xiang T, Liang B, Deng H, Wang H, Feng X, et al. Characterization of SARS-CoV-2-Specific Humoral and Cellular Immune Responses Induced by Inactivated COVID-19 Vaccines in a Real-World Setting. Frontiers in immunology. 2021;12:802858. PubMed PMID: 35003131. Pubmed Central PMCID: 8727357.

12. Watson OJ, Barnsley G, Toor J, Hogan AB, Winskill P, Ghani AC. Global impact of the first year of COVID-19 vaccination: a mathematical modelling study. The Lancet Infectious diseases. 2022 Sep;22(9): 1293–302. PubMed PMID: 35753318. Pubmed Central PMCID: 9225255 GlaxoSmithKline, and WHO related to COVID-19 epidemiology and from The Global Fund to Fight AIDS, Tuberculosis and Malaria for work unrelated to COVID-19. ACG is a non-remunerated member of scientific advisory boards for Moderna and the Coalition for Epidemic Preparedness. ABH and PW have received personal consultancy related to COVID-19 work from WHO. All other authors declare no competing interests.

13. Li Y, Wang X, Jin J, Ma Z, Liu Y, Zhang X, et al. T-cell responses to SARS-CoV-2 Omicron spike epitopes with mutations after the third booster dose of an inactivated vaccine. Journal of medical virology. 2022 Aug;94(8):3998–4004. PubMed PMID: 35474581. Pubmed Central PMCID: 9088599.

14. Cucunawangsih C, Wijaya RS, Lugito NPH, Suriapranata I. Antibody response after a third dose mRNA-1273 vaccine among vaccinated healthcare workers with two doses of inactivated SARS-CoV-2 vaccine. International journal of infectious diseases: IJID: official publication of the International Society for Infectious Diseases. 2022 May;118:116–8. PubMed PMID: 35192955. Pubmed Central PMCID: 8857755.

15. Mok CKP, Cohen CA, Cheng SMS, Chen C, Kwok KO, Yiu K, et al. Comparison of the immunogenicity of BNT162b2 and CoronaVac COVID-19 vaccines in Hong Kong. Respirology. 2022 Apr;27(4):301–10. PubMed PMID: 34820940. Pubmed Central PMCID: 8934254.

16. van Doremalen N, Lambe T, Spencer A, Belij-Rammerstorfer S, Purushotham JN, Port JR, et al. Publisher Correction: ChAdOx1 nCoV-19 vaccine prevents SARS-CoV-2 pneumonia in rhesus macaques. Nature. 2021 Feb;590(7844):E24. PubMed PMID: 33469217.

17. Schultz BM, Melo-Gonzalez F, Duarte LF, Galvez NM, Pacheco GA, Soto JA, et al. A booster dose of an inactivated SARS-CoV-2 vaccine increases neutralizing antibodies and T cells that recognize Delta and Omicron variants of concern. medRxiv: the preprint server for health sciences. 2022 Feb 7. PubMed PMID: 35441179. Pubmed Central PMCID: 9016658.

18. Gao Y, Cai C, Grifoni A, Muller TR, Niessl J, Olofsson A, et al. Ancestral SARS-CoV-2-specific T cells cross-recognize the Omicron variant. Nature medicine. 2022 Mar;28(3):472–6. PubMed PMID: 35042228. Pubmed Central PMCID: 8938268.

19. Willems LH, Nagy M, Ten Cate H, Spronk HMH, Jacobs LMC, Kranendonk J, et al. ChAdOx1 vaccination, blood coagulation, and inflammation: No effect on coagulation but increased interleukin-6. Research and practice in thrombosis and haemostasis. 2021 Dec;5(8):e12630. PubMed PMID: 34934894. Pubmed Central PMCID: 8652129.

20. Farsakoglu Y, Palomino-Segura M, Latino I, Zanaga S, Chatziandreou N, Pizzagalli DU, et al. Influenza Vaccination Induces NK-Cell-Mediated Type-II IFN Response that Regulates Humoral Immunity in an IL-6-Dependent Manner. Cell reports. 2019 Feb 26;26(9):2307–15 e5. PubMed PMID: 30811982.

21. Barrett JR, Belij-Rammerstorfer S, Dold C, Ewer KJ, Folegatti PM, Gilbride C, et al. Phase 1/2 trial of SARS-CoV-2 vaccine ChAdOx1 nCoV-19 with a booster dose induces multifunctional antibody responses. Nature medicine. 2021 Feb;27(2):279–88. PubMed PMID: 33335322.

22. Clem AS. Fundamentals of vaccine immunology. Journal of global infectious diseases. 2011 Jan;3(1):73–8. PubMed PMID: 21572612. Pubmed Central PMCID: 3068582.

23. Laidlaw BJ, Ellebedy AH. The germinal centre B cell response to SARS-CoV-2. Nature reviews Immunology. 2022 Jan;22(1):7–18. PubMed PMID: 34873279. Pubmed Central PMCID: 8647067.

24. Stavnezer J, Schrader CE. IgH chain class switch recombination: mechanism and regulation. Journal of immunology. 2014 Dec 1;193(11):5370–8. PubMed PMID: 25411432. Pubmed Central PMCID: 4447316.

25. Mesin L, Ersching J, Victora GD. Germinal Center B Cell Dynamics. Immunity. 2016 Sep 20;45(3):471–82. PubMed PMID: 27653600. Pubmed Central PMCID: 5123673.

26. Cheng ZJ, Huang H, Liu Q, Zhong R, Liang Z, Xue M, et al. Immunoassay and mass cytometry revealed immunological profiles induced by inactivated BBIBP COVID-19 vaccine. Journal of medical virology. 2022 Nov;94(11):5206–16. PubMed PMID: 35801663. Pubmed Central PMCID: 9350407.

27. Vadrevu KM, Ganneru B, Reddy S, Jogdand H, Raju D, Sapkal G, et al. Persistence of immunity and impact of third dose of inactivated COVID-19 vaccine against emerging variants. Scientific reports. 2022 Jul 14;12(1):12038. PubMed PMID: 35835822. Pubmed Central PMCID: 9281359.

